# Fatty Acid Transport Protein-2 (FATP2) Inhibition Enhances Glucose Tolerance through α-Cell-mediated GLP-1 Secretion

**DOI:** 10.1101/2025.01.31.635976

**Authors:** Shenaz Khan, Robert J. Gaivin, Zhiyu Liu, Vincent Li, Ivy Samuels, Jinsook Son, Patrick Osei-Owusu, Jeffrey L. Garvin, Domenico Accili, Jeffrey R. Schelling

## Abstract

Type 2 diabetes affects more than 30 million people in the US, and a major complication is kidney disease. During the analysis of lipotoxicity in diabetic kidney disease, global fatty acid transport protein-2 (FATP2) gene deletion was noted to markedly reduce plasma glucose in db/db mice due to sustained insulin secretion. To identify the mechanism, we observed that islet FATP2 expression was restricted to α-cells, and α-cell FATP2 was functional. Direct evidence of FATP2KO-induced α-cell-mediated GLP-1 secretion included increased GLP-1-positive α-cell mass in FATP2KO db/db mice, small molecule FATP2 inhibitor enhancement of GLP-1 secretion in αTC1-6 cells and human islets, and exendin[9-39]-inhibitable insulin secretion in FATP2 inhibitor-treated human islets. FATP2-dependent enteroendocrine GLP-1 secretion was excluded by demonstration of similar glucose tolerance and plasma GLP-1 concentrations in db/db FATP2KO mice following oral versus intraperitoneal glucose loading, non-overlapping FATP2 and preproglucagon mRNA expression, and lack of FATP2/GLP-1 co-immunolocalization in intestine. We conclude that FATP2 deletion or inhibition exerts glucose-lowering effects through α-cell-mediated GLP-1 secretion and paracrine β-cell insulin release.

**Graphical abstract:** 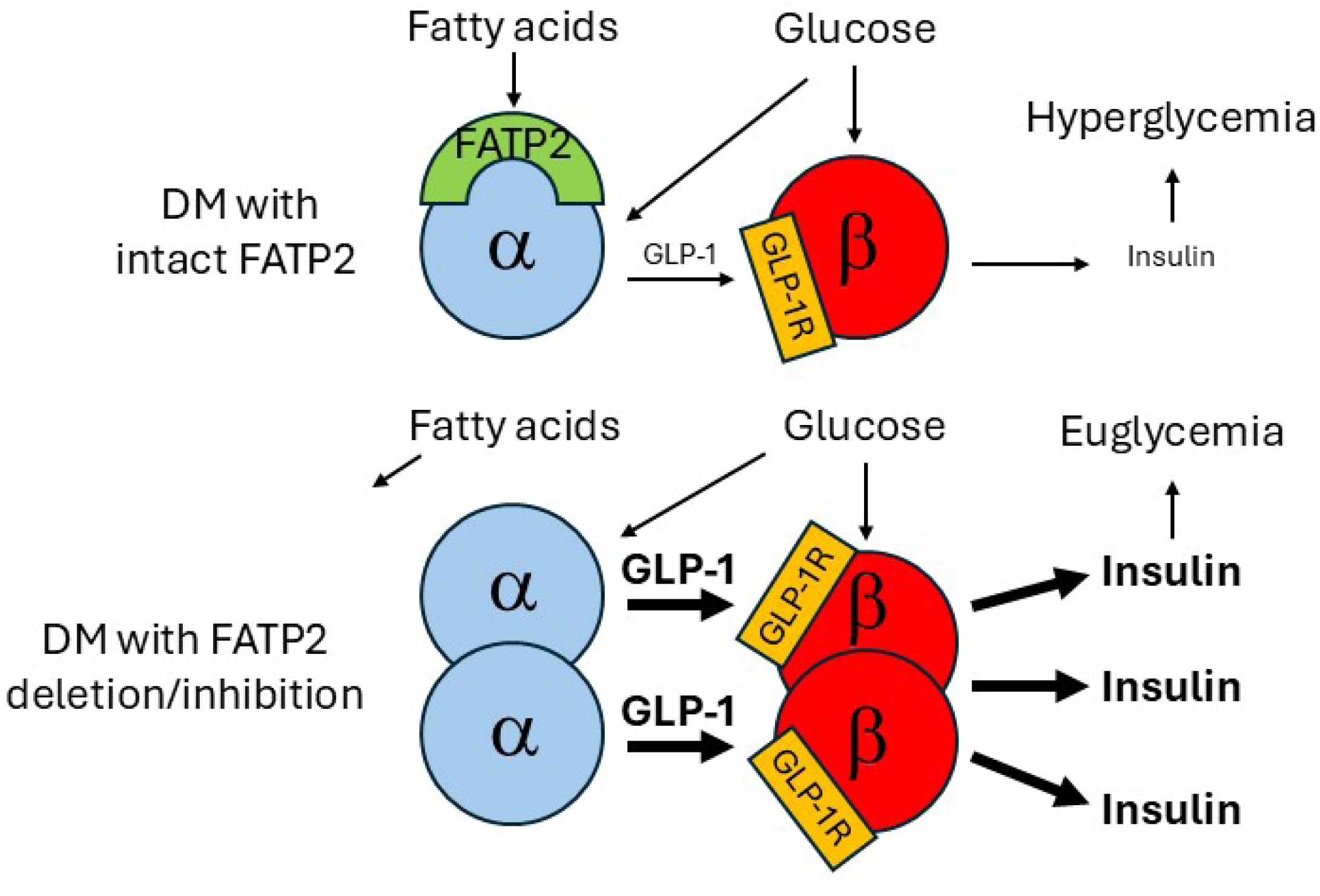

## INTRODUCTION

Type 2 diabetes affects more than 30 million people in the US (830 million worldwide) and is a major public health problem due to morbidity and mortality from microvascular (kidney disease, retinopathy, neuropathy) and macrovascular (myocardial infarction, stroke, peripheral vascular) complications. The pathophysiologic mechanisms are complex, with significant contributions from altered glucose and lipid metabolism (1, 2).

The lipid abnormalities in diabetes include increased plasma fatty acid concentrations (3). Fatty acids circulate predominantly as non-covalently bound complexes with albumin or as covalently-linked esters with glycerol to form triglycerides. Cellular fatty acid uptake is facilitated by a family of six evolutionarily conserved plasma membrane fatty acid transport proteins (FATP1-6), which are expressed in a tissue-specific fashion (4).

FATP2 is a major fatty acid transporter in liver and kidney, and has been implicated in the pathophysiology of metabolic dysfunction-associated steatotic liver disease (MASLD) and diabetic kidney disease (5). Modest reduction of fasting plasma glucose was observed in mice with liver-specific FATP2 gene (*Slc27a2*) deletion (6). Inhibition of FATP2 by shRNA tail vein injection in high fat diet-induced diabetic mice also resulted in mild plasma glucose reduction, as well as improved insulin sensitivity (7). Because tail vein-injected siRNA uptake is primarily by liver (8), it was assumed that the hypoglycemic effect of FATP2 inhibition was mediated by enhanced hepatic glucose metabolism. However, tail vein-injected reporter siRNAs are detectable in other FATP2-expressing organs, including intestine and pancreas (9–11), which raises the possibility that extrahepatic FATP2 inhibition contributes to the glucose-lowering phenotype.

In contrast to the modest glucose reduction with liver FATP2 inhibition (6, 7), global FATP2 deletion was associated with profoundly lower plasma glucose in genetic and inducible mouse models of type 2 diabetes (9). Furthermore, diabetic mice with intact FATP2 developed islet atrophy and reduced plasma insulin, whereas diabetic mice with FATP2 deletion demonstrated islet hypertrophy and sustained hyperinsulinemia (9). These observations suggest that FATP2 inhibition enhances pancreatic β-cell mass and function. Localization of FATP2 to specific pancreatic cells, and assignment of FATP2 inhibition to pancreatic endocrine functions, have not been previously described.

Glucose-stimulated insulin secretion (GSIS) is augmented by glucagon-like peptide (GLP)-1 in diabetes (12). Following a glucose-or fat-containing meal, GLP-1 is secreted into the circulation by enteroendocrine L-cells in the distal ileum and proximal colon, and ultimately binds to GLP-1 receptors on pancreatic β-cells to stimulate insulin secretion (13). Fatty acids can also directly stimulate GSIS through binding to free fatty acid receptor 1 (FFAR1), which is expressed at low levels on β-cells (14, 15). The insulinotropic effects of fatty acids in acute models are counterbalanced by the possibility of chronic fatty acid-induced α-and/or β-cell lipoapoptosis (16), which may be mediated by specific fatty acid receptors.

Pancreatic α-cells also secrete GLP-1, which enhances GSIS through paracrine activation of the β-cell GLP-1 receptor (12, 15, 17), particularly under conditions of β-cell stress (18). The primacy of paracrine GLP-1 is supported by observations that GLP-1 secreted by enteroendocrine cells has a half-life of only two minutes (19), due to proteolysis by local dipeptidyl peptidase (DPP)-4. It has therefore been postulated that gut-derived GLP-1 may not achieve sufficient concentration to stimulate distant β-cell GLP-1 receptors, and an α-cell, rather than enteroendocrine source of GLP-1, regulates insulin secretion under diabetic conditions (18).

With the emergence of GLP-1 receptor agonists as weight loss drugs, there is intense recent interest regarding GLP-1 regulation of lipid metabolism. However, the effect of fatty acids on GLP-1 biology is much less well understood, and the influence of FATP2 inhibition on GLP-1 pathways has not been investigated. In this report we describe mechanisms of insulinotropic activity by FATP2 inhibition, through augmentation of α-cell-mediated GLP-1 secretion.

## RESEARCH DESIGN AND METHODS

### Mice

db/db and db/db FATP2 KO mice on a C57BL-KS/J congenic background have previously been described (9). Conditional proximal tubule FATP2KO mice were generated from intercrosses between GGT1-Cre (JaxLabs) and Slc27a2-floxed mice [gift from Dr. Dmitry Gabrilovich (20)] on a congenic C57BL-KS/J background. Genotyping by PCR from toe samples was done by Transnetyx. Type 2 diabetes was induced as previously described (9). Briefly, at six weeks of age, male mice were fed a high fat diet (Harlan, Teklad TD.06414, 60.3% fat, 21.3% carbohydrate, 18.4% protein) for six months. After three months of high fat diet, mice were administered low dose intraperitoneal streptozotocin (45 μg/g) daily for three consecutive days. Diabetes was defined by fasting glucose greater than 200 mg/dL. Glucose and GLP-1 were assayed after fasting from 6:00-10:00 AM. Tail vein blood glucose was assayed by glucometer, as previously described (9). All studies were conducted in accordance with protocols approved by the Institutional Animal Care and Use Committee of Case Western Reserve University School of Medicine.

### Immunohistochemistry

Paraffin-embedded human ileum and duodenum slides were purchased from Zyagen. Mouse pancreas samples were fixed in paraformaldehyde (4%, 24 hrs, room temp), cryopreserved in sucrose (30% in PBS overnight) and frozen at-80° C. Frozen sections (5 µm by cryostat) were permeabilized with Triton X-100 (Millipore Sigma; 0.2% in PBS, 10 minutes, room temp) and blocked with donkey serum (5% in PBS, 1 hour, room temp).

Primary antibodies are listed in Supplementary Table 1. Sections were mounted in SlowFade Diamond Antifade Mountant with DAPI (Invitrogen) and viewed with a Leica or Olympus confocal microscope.

### β-cell and GLP-1-positive α-cell mass

α- and β-cell mass were calculated using published methods (21). Mouse pancreas was fixed in paraformaldehyde (4%, 24 hrs, room temp), weighed, cryopreserved in sucrose (30% in PBS overnight) and frozen at-80° C. Three 5 µm frozen sections, cut 200 µm apart, were analyzed. α- and β-cells were labeled with glucagon and insulin antibodies, respectively, as previously described (9). α- and β-cell areas were normalized to total pancreatic area, and these values were then multiplied by pancreas weight to obtain α- and β-cell mass. GLP-1-positive α-cell mass was determined by multiplying the α-cell mass and the percentage of glucagon-positive cells that co-labeled with GLP-1 antibodies.

### Reverse transcriptase polymerase chain reaction (RT-PCR)

Methods have previously been described in detail (9). Briefly, total RNA was extracted from whole mouse organs, αTC1-6 (ATCC) or INS-1 (gift from Dr. Yisheng Yang, Case Western Reserve University) cells using RNeasy Mini Kit (Qiagen). RNA concentrations were determined using the NanoDrop 2000 Spectrophotometer (Thermo Scientific). RT was performed using 5 µg total RNA, and cDNA was generated using SuperScript III First-Strand Synthesis System (Invitrogen). Human α-cell cDNA was purchased from Celprogen. PCR reactions from 1.5 µg cDNA were conducted in 20 µl volume using EmeraldAmp Max PCR Master Mix 2X premix (Takara Bio Inc.), according to recommended protocol and PCR cycling conditions, for 30 cycles (35 cycles for INS-1 cell PCR). Primers were purchased from Eurofins Genomics and sequences are shown in Supplementary Table 2. PCR products underwent 2% agarose gel electrophoresis and bands were identified by ethidium bromide (Invitrogen) staining and photographed. Quantitative PCR (qPCR) was conducted as previously described (22). Briefly, cDNA was generated using SuperScript III First-Strand, and amplified using the Radiant SYBR Green 2x Master Mix (Alkali Scientific) and QuantStudio 3 System (Applied Biosystems). Quantification was determined by the comparative C_T_ (ΔΔC_T_) method.

### FA uptake in αTC1-6 cells

Experiments were conducted according to previously described methods (23). Briefly, αTC1-6 cells were seeded in 96-well plates, and cultured to confluence over 24 h. Wells were washed with serum-free, phenol-free media for 2 h at 37° C; Lipofermata (MedChemExpress) or compound 5668437 (Chembridge) were robotically incubated for the final hour. BODIPY-conjugated C18 fatty acids (Molecular Devices QBT assay, 2.5 μM complexed with 0.2 % fatty acid-free albumin carrier in PBS + FATP2 inhibitors) were robotically added at time = 0. Excitation λ = 490 nm pulses were delivered, and emission λ = 510 nm was recorded at 15-sec intervals for 10 min. BODIPY-labeled fatty acid uptake velocity was determined from 2 min fluorescence values. Plates were imaged on the Synergy Neo2 HTX Multi-Mode Microplate reader (BioTek) and averaged from six fields captured from each well using Gen5 software. IC_50_ values were calculated using GraphPad Prism 7 software.

### Hormone assays

Glucagon (10-I271-01), GLP-1 (10-I278-01), and human insulin (10-1113-01) were assayed from mouse plasma or culture media by ELISA (Mercodia).

### In vivo metabolic analyses

To facilitate multiple blood samples for GLP-1 assays, a carotid artery catheter was placed the day prior to experiments. Mice were fasted for four hours (6:00 AM-10:00 AM) prior to oral (OGTT, 2 g/kg by gavage) or intraperitoneal (IPGTT, 2 g/kg) glucose tolerance tests (24). Arterial blood was drawn at baseline and then one hr after glucose administration, and plasma was saved at-80° C in tubes containing linagliptin (MedChemExpress; 100 nM) for GLP-1 assays later. Glucometer readings were obtained at baseline, 30 min, 60 min, 90 min and 120 min. Non-fasting blood samples for glucagon were obtained in mice by cardiac puncture at the time of sacrifice.

### Glucose-stimulated GLP-1 and insulin secretion

Human islets (ProdoLabs) were cultured according to established methods (25). Islets were identified under a dissecting microscope and suspended in RPMI 1640 + 15% FBS for at least 16 hr prior to experimentation. Islets were equilibrated in petri dishes containing modified Krebs buffer (2 mM NaHCO_3_, 10 mM HEPES, pH 7.4, 37° C, 1 hr). Ten islets were selected and deposited in cell culture wells, with quintuplicate wells for each condition. Initial incubations included modified Krebs buffer supplemented with 2.8 mM glucose (37° C, 1 hr) plus linagliptin (MedChemExpress; 100 nM) ± palmitate (Avanti; 100-400 µM complexed with 0.2% delipidated albumin), Lipofermata (MedChemExpress; 50 µM), or exendin[9-39] (MedChemExpress; 100 nM). After 1 hour, minimum volumes of media for GLP-1 or insulin ELISA assays in duplicate were saved at-80° C. Identical conditions were then repeated for an additional hour in Krebs buffer with high glucose (16.8 mM, 37° C, one hour). Islets were pelleted by centrifugation, lysed in SDS-PAGE buffer, and assayed for protein content (Nanodrop; absorption at λ = 280 nm). A similar protocol was followed for glucose-stimulated GLP-1 secretion in αTC1-6 cells, except for incubations in 5 mM and 25 mM glucose, to conform with established methods for these cells (26, 27).

### Statistics

Graphical data are presented as mean ± SEM and analyzed using GraphPad Prism 7 software. Data from multiple groups were analyzed by one-way ANOVA and Tukey’s post-hoc test for multiple comparisons. Data from two groups were analyzed by unpaired two-tailed *t* test. Statistical significance for all analyses is defined as a *P* value <0.05.

## RESULTS

Diabetic mice with global FATP2 gene deletion (FATP2KO) developed markedly reduced fasting plasma glucose (9). FATP2 is most abundantly expressed in kidney, and within kidney, exclusively in the apical proximal tubule membrane (23, 28). The proximal tubule contributes to gluconeogenesis, particularly in the pathogenesis of diabetes (29). However, deletion of proximal tubule FATP2 in an inducible model of diabetes did not alter fasting plasma glucose concentrations (Supplementary Figure 1). These data suggest that the glucose-lowering effect in global FATP2KO mice is not due proximal tubule FATP2 gene deletion, but rather by an extrarenal mechanism.

Pancreatic islet FATP2 protein expression is highly upregulated in the setting of elevated glucose concentration (30), and global FATP2 gene deletion in diabetic mice was associated with increased islet area and sustained plasma insulin (9), suggesting that induction of FATP2 on islet cells mediates pancreatic islet dysfunction. Compared to diabetic mice with intact FATP2, diabetic *Lepr^db/db^*(db/db) mice with FATP2KO demonstrated islet hypertrophy (Figures 1A and 1B, respectively) and increased β-cell mass (Figure 1C). These data are consistent with FATP2 deletion causing rescue of β-cell failure in db/db mice (9, 31).

**Figure 1.**
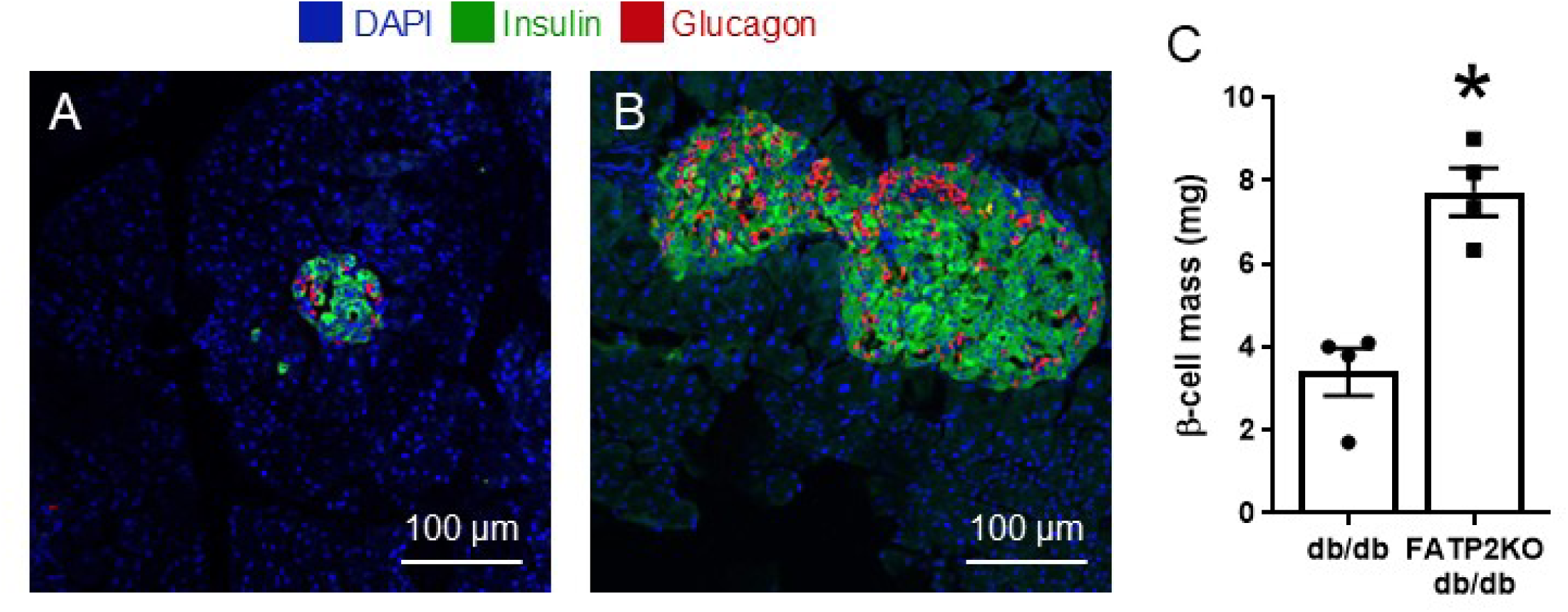
I**slet hypertrophy and increased β-cell mass in FAT2KO db/db mice.** Representative immunohistochemistry images of pancreatic islets from db/db (**A**) and FATP2KO db/db (**B**) mice. α-and β-cells were labeled as described in Methods. (**C**) β-cell mass was calculated as described in Methods for db/db and FATP2KO db/db mice.

FATP2 mRNA is expressed in pancreas (9), and variably in α-and β-cells from scRNAseq databases (32–37). Figure 2 demonstrates that FATP2 protein co-localized exclusively with α-, but not β- or δ-cells. FATP2 mRNA is expressed in mouse and human pancreas tissue and α-cells (Figures 3A and 3B), but is undetectable in INS-1 β-cells (not shown). To determine whether α- cell FATP2 is functional, long-chain fatty acid transport was measured in αTC1-6 cells. Fatty acid uptake was inhibited by the FATP2 inhibitor, Lipofermata (Figure 4, IC_50_ = 5.4 µM), and related compound 5668437 (Supplementary Figure 2, IC_50_ = 0.8 µM). The IC_50_ values for both inhibitors are in agreement with other epithelial cells (38, 39). The conclusion from these experiments is that α-cells express functional FATP2, whereas β-cells do not.

**Figure 2.**
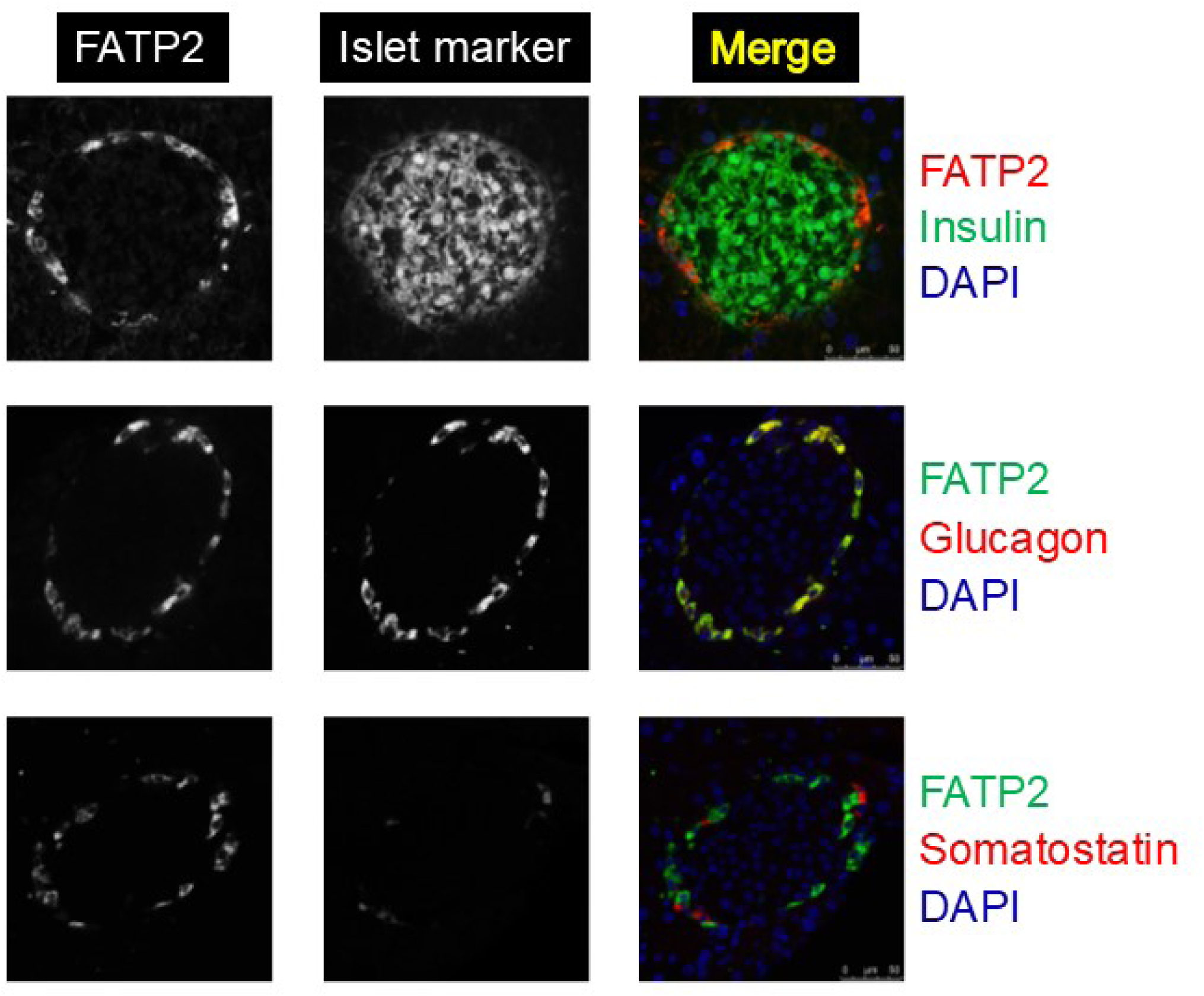
FATP2 expression localizes to pancreatic α-cells. Wild-type mouse pancreas islets were immunohistochemically labeled for FATP2 expression in α-, β- and δ-cells as described in Methods. Merged images, representing cell-specific FATP2 expression are shown in yellow.

**Figure 3.**
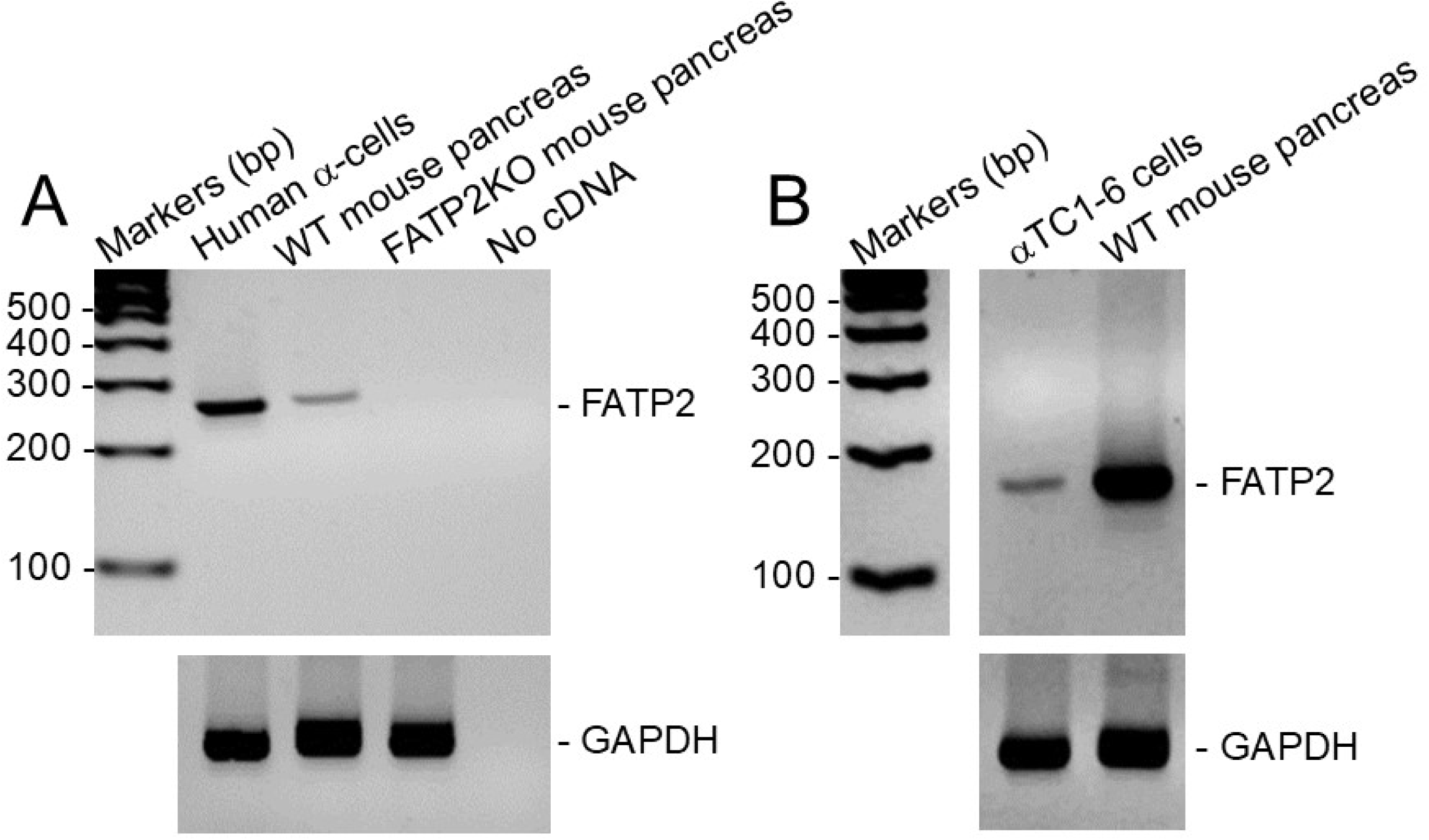
FATP2 mRNA is expressed in pancreatic α-cells. FATP2 and loading control GAPDH mRNA expression were determined in human and mouse pancreatic tissue and α-cell lines by RT-PCR (as described in Methods).

**Figure 4.**
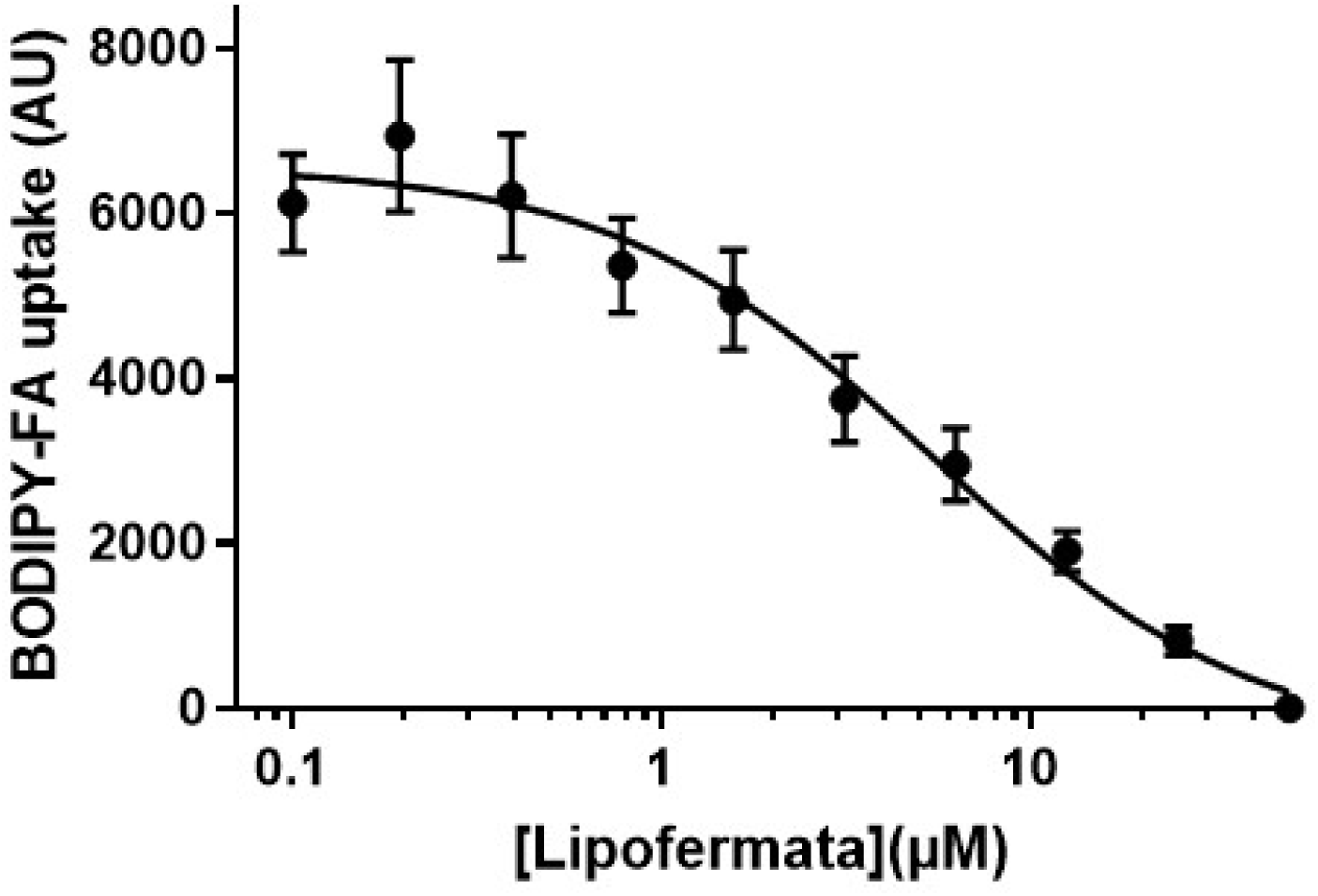
FATP2-dependent fatty acid uptake in αTC1-6 cells. Mouse αTC1-6 cells were preincubated with Lipofermata (1 hr, 37° C, 0–50 μM) in triplicate. BODIPY-labeled fatty acid uptake velocity was then determined, as described in Methods. Results are from four experiments.

We next focused on the mechanism by which α-cell FATP2 deletion regulates insulin secretion. GLP-1 and glucagon bind with high and low affinity, respectively, to the GLP-1 receptor on the β-cell, which facilitates GSIS (12). Random (non-fasting) plasma glucagon levels were increased in db/db mice with intact FATP2. Plasma glucagon concentrations were decreased in FATP2KO db/db mice, and not significantly different compared to wild-type (Supplementary Figure 3).

The relative contribution of enteroendocrine L-cell-versus α-cell-derived GLP-1 on β-cell insulin secretion has been debated (40). To investigate whether enteroendocrine cells are the GLP-1 source in FATP2KO mice, OGTT and IPGTT were conducted in db/db mice with or without FATP2 gene deletion. The rationale is if the major GLP-1 source is enteroendocrine, oral glucose would stimulate a greater increase in plasma GLP-1 and superior glucose tolerance compared to IP glucose (41). Four-month-old db/db and FATP2KO db/db mice were obese, though baseline weights (45 ± 9 g and 54 ± 10 g, respectively), were similar (P >0.05). Figure 5A shows markedly lower fasting plasma glucose concentrations in FATP2KO db/db compared to db/db mice, consistent with previous reports (9). Figures 5A and 5B show no difference between OGTT and IPGTT in FATP2KO db/db mice. Glucose disposal was also similar following OGTT vs. IPGTT in non-diabetic FATP2KO mice (Supplementary Figure 4A), as well as FATP2KO db/db *eNOS^-/-^* mice (Supplementary Figure 4B), which phenocopy diabetic kidney disease (9, 42, 43). Importantly, plasma GLP-1 increases were similar in FATP2KO db/db mice after oral and IP glucose loading (Figure 5C). The lack of enhanced glucose tolerance and GLP-1 concentration with oral glucose suggests that α-cells, rather than enteroendocrine L-cells, are the source of GLP-1 in mice with global FATP2 gene deletion.

**Figure 5.**
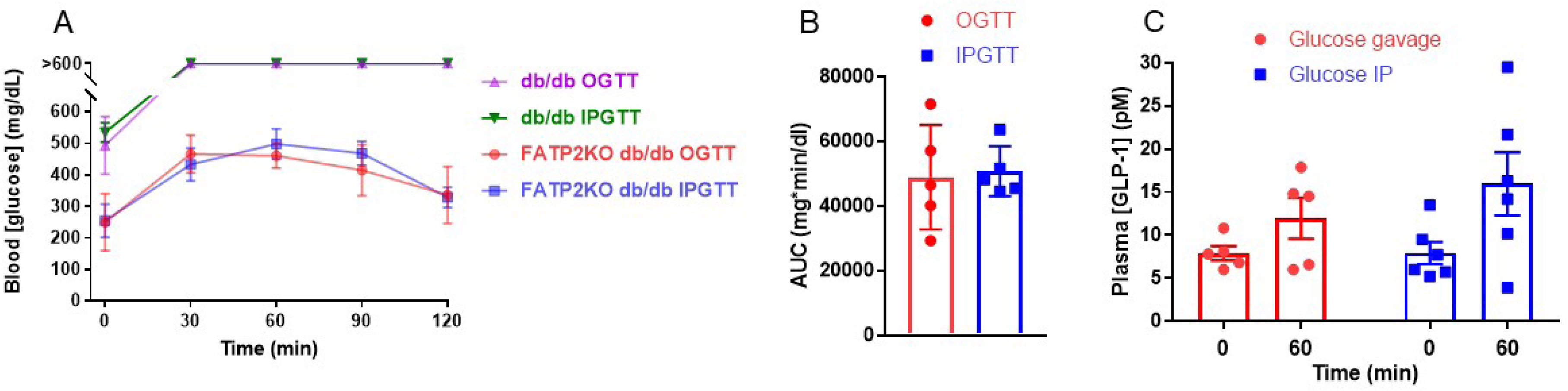
Blood glucose and plasma GLP-1 concentrations following oral (OGTT) and intraperitoneal (IPGTT) glucose tolerance tests. (**A**) OGTT and IPGTT were conducted in db/db and FATP2KO db/db mice, as described in Methods. Blood glucose was determined at the indicated times in five mice per group. (**B**) As an index of glucose disposal, the area under the curve (AUC) corresponding to FATP2KO db/db experiments in (**A**) was integrated using GraphPad Prism 7 software. (**C**) Plasma was obtained at baseline and at the one hr time point during OGTT or IPTGG in FATP2KO db/db mice.

To further address the possibility of FATP2 effects on enteroendocrine GLP-1 secretion, FATP2 and GCG (preproglucagon gene encoding GLP-1) mRNA expression was evaluated in intestine segments by qPCR. Both transcripts were detected throughout the mouse GI tract, but with distinct patterns, and minimal overlap (Figure 6A and 6B). Protein expression by immunohistochemistry in human distal ileum (Figure 6C and 6D) and duodenum (Figure 6E and 6F) demonstrated no FATP2/GLP-1 co-localization. Taken together, the data suggest that FATP2 deletion does not directly influence GLP-1 release by enteroendocrine cells.

**Figure 6.**
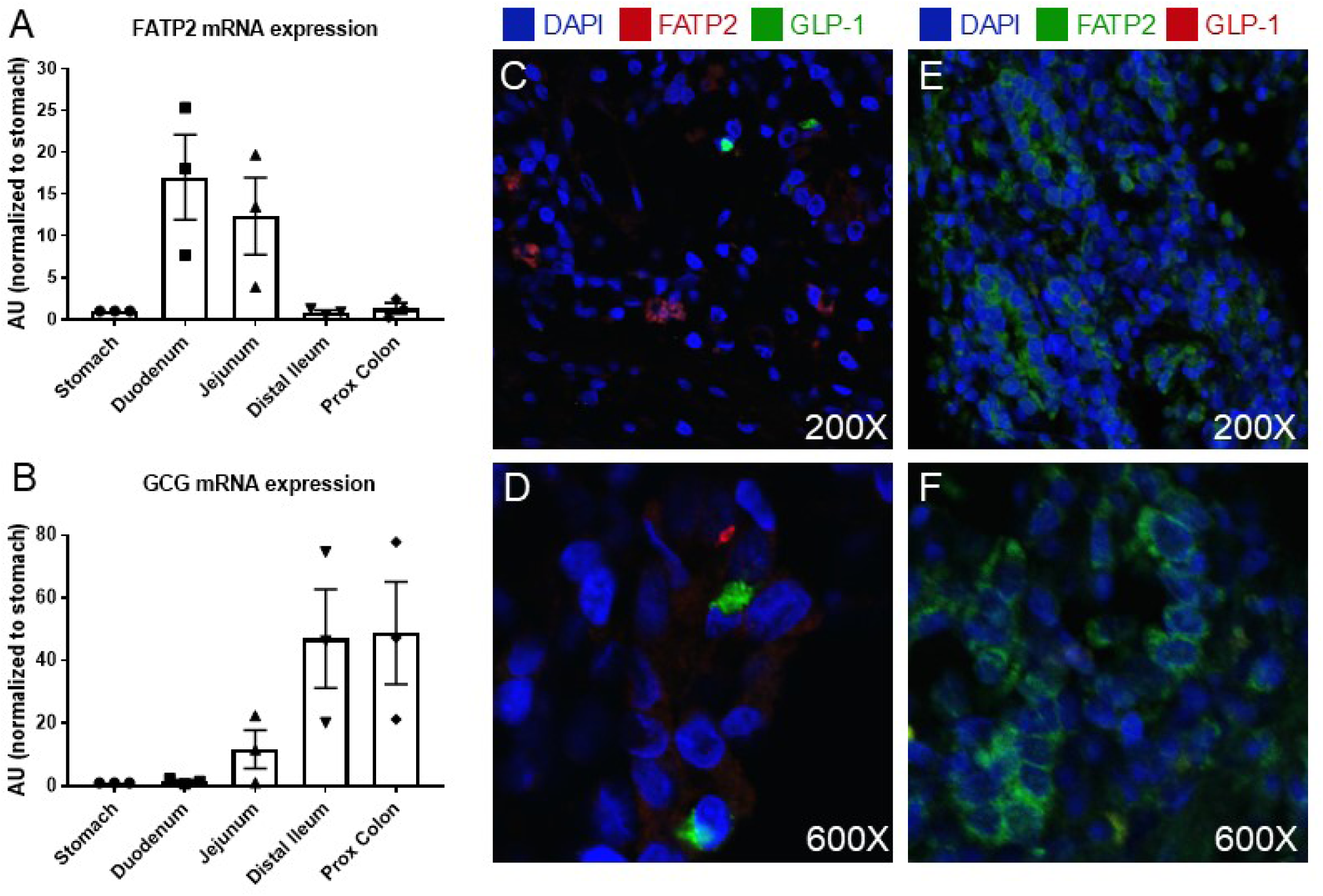
FATP2 and GLP-1 localization in intestine. FATP2 (**A**) and preproglucagon (**B**) mRNA expression were determined in mouse gut segments by qPCR, as described in Methods. Data are normalized to stomach expression, which is defined as 1.0. Immunohistochemical labeling of FATP2 and GLP-1 in human distal ileum (**C** and **D**) and duodenum (**E** and **F**).

The next set of experiments tested the direct effects of FATP2 inhibition on α-cell GLP-1 secretion. Both α-cell mass and % of GLP-1-positive α-cells were increased in FATP2KO db/db islets (Supplementary Figure 5). Consequently, the product of α-cell mass and % of GLP-1-positive α-cells (GLP-1-positive α-cell mass) was significantly greater in FATP2KO db/db compared to db/db mice (Figure 7A). To determine whether α-cell GLP-1 secretion is regulated by FATP2 inhibition, the effect of FATP2 inhibition on glucose-stimulated GLP-1 secretion was measured in human islets. Figure 7B shows that the FATP2 inhibitor Lipofermata enhanced glucose-stimulated GLP-1 secretion, particularly under high glucose conditions. Prolonged high glucose plus palmitate has previously been shown to inhibit GLP-1 secretion, due to glucolipotoxicity (44). To assess whether FATP2 inhibition preserves GLP-1 secretion through prevention of glucolipotoxicity, glucose-stimulated GLP-1 release was tested in αTC1-6 cells (26, 27), in response to palmitate with or without Lipofermata preincubation (38, 39). Figure 7C demonstrates that under low and high glucose conditions, palmitate decreased GLP-1 secretion, which was rescued by Lipofermata preincubation. Taken together, the data indicate that FATP2 inhibition or deletion preserves α-cell GLP-1 secretion.

**Figure 7.**
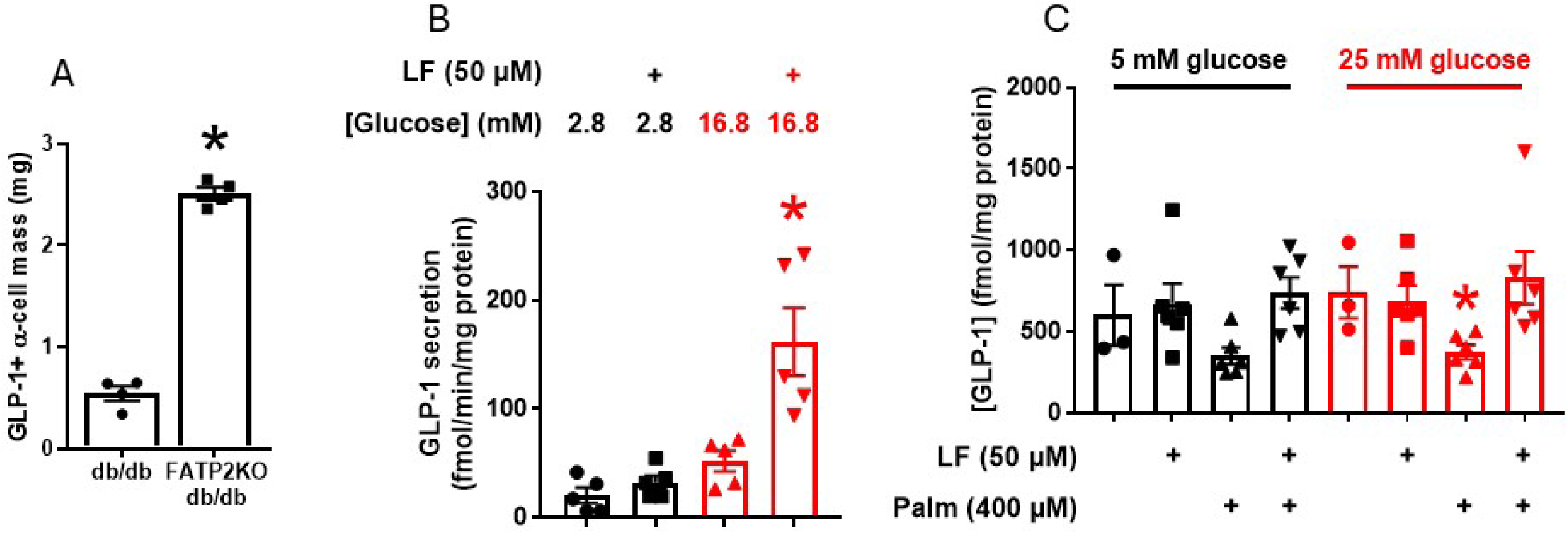
FATP2KO/deletion effect on glucose-stimulated GLP-1. (**A**) Pancreas GLP-1-positive α- cell mass was determined as described in Methods in db/db and FATP2KO db/db mice. * P <0.01 compared to db/db group. (**B**) Human islets were pre-incubated with or without Lipofermata (LF), then tested for glucose-stimulated GLP-1 secretion as described in Methods. * P <0.01 compared to all other groups. (**C**) αTC1-6 cells were pe-incubated with or without Lipofermata (LF) or palmitate (Palm) as indicated. Glucose-stimulated GLP-1 secretion was then measured as described in Methods. * P <0.05 compared LF + Palm.

To investigate whether FATP2 inhibition regulates α-cell GLP-1-dependent insulin secretion, GSIS was examined in human islets pretreated with palmitate, Lipofermata and/or the GLP-1 receptor inhibitor exendin[9-39]. Figure 8 demonstrates that Lipofermata enhanced insulin secretion (particularly under high glucose concentration conditions). Importantly, a large proportion of the increase was exendin[9-39]-inhibitable, indicating that FATP2 inhibition enhanced α-cell secretion of GLP-1, which acts in a paracrine manner to enhance GSIS.

**Figure 8.**
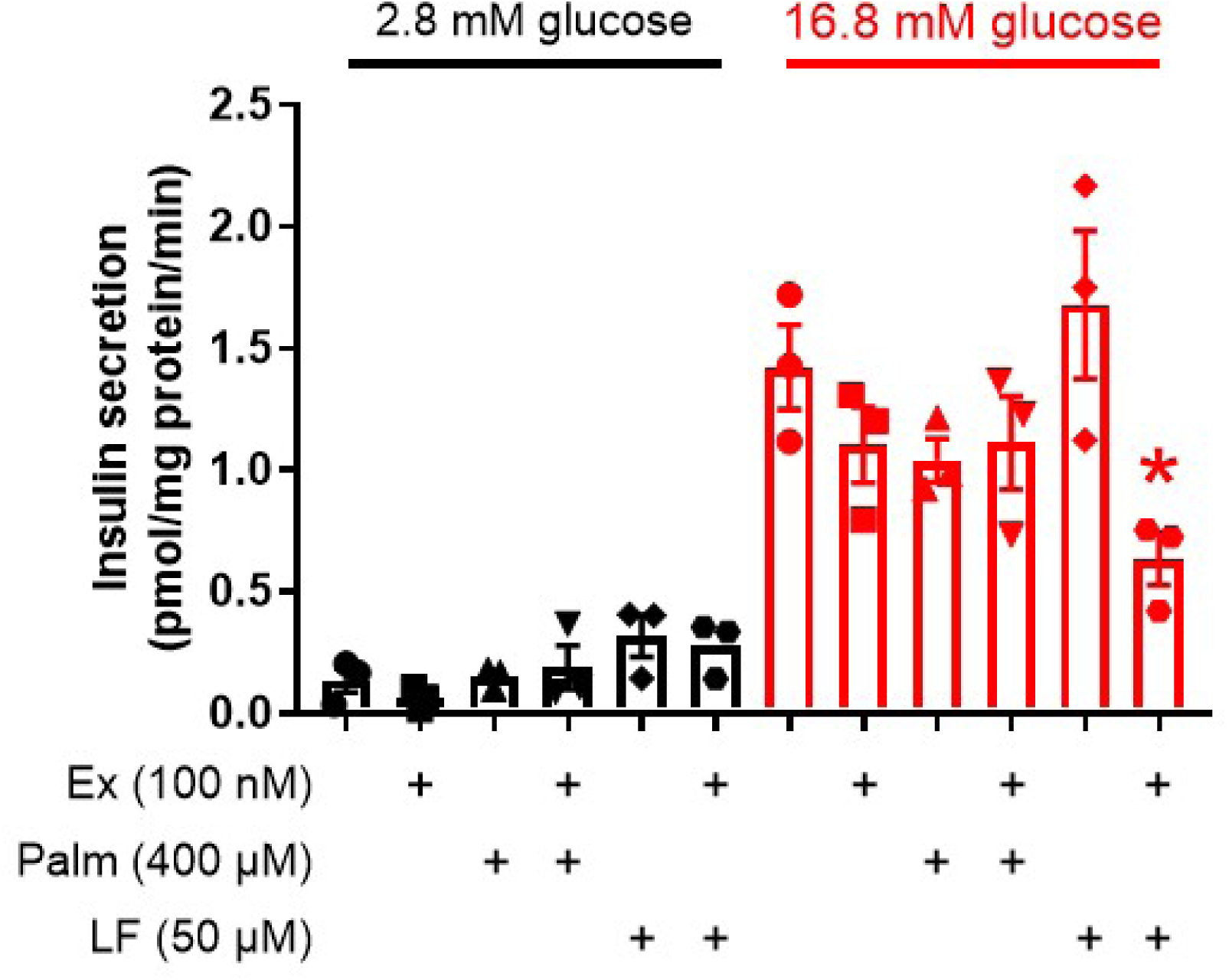
Effect of FATP2 inhibition on GLP-1-stimulated insulin secretion in human islets. Glucose-stimulated insulin secretion was measured in human islets, which were pre-incubated with Lipofermata (LF) and then exposed to exendin[9-39] (Ex) or palmitate (Palm). * P <0.01 compared to 16.8 mM glucose + LF group.

## DISCUSSION

Type 2 diabetes is characterized by initial hyperinsulinemia, and subsequent β-cell exhaustion and insulin deficiency. We previously showed that global FATP2 deletion in genetic and inducible mouse models of type 2 diabetes was associated with markedly decreased plasma glucose and increased plasma insulin (9). Using in vivo, ex vivo and in vitro models, we now show that the mechanism of sustained hyperinsulinemia in the setting of FATP2 inhibition or deletion is α-cell-mediated GLP-1 secretion, with paracrine stimulation of β-cell insulin secretion (graphical abstract).

The conventional dogma is that intestinal L-cells are the major source of GLP-1, which exerts potent insulinotropic effects. This mechanism has been questioned, though, due to the short half-life, and potentially insufficient GLP-1 concentration to stimulate GLP-1 receptors on distant β-cells. We provide substantial evidence against an enteroendocrine source of GLP-1 as the mechanism for increased plasma insulin in FATP2 KO mice, including (a) no difference between plasma GLP-1 or glucose tolerance in db/db FATP2KO mice following oral vs. IP glucose loading, (b) non-overlapping FATP2 and GCG mRNA expression in intestine, and (c) lack of FATP2 and GLP-1 protein co-localization in intestinal cells. Additional evidence against FATP2KO regulation of enteroendocrine GLP-1 is the lack of weight loss in diabetic FATP2KO mice. The mechanism of GLP-1-mediated weight reduction is complex, but at least partly involves gut-derived GLP-1 stimulation of the vagus nerve, which leads to anorexia and delayed gastric emptying (45). FATP2KO db/db mice were slightly heavier than db/db mice, as were FATP2KO db/db eNOS^-/-^ compared to db/db eNOS^-/-^ mice (9), which mitigates against an enteroendocrine GLP-1 mechanism. Furthermore, the lack of weight loss in FATP2KO db/db eNOS^-/-^ mice was accompanied by no difference in food intake compared to db/db eNOS^-/-^ mice (9), which argues against a FATPKO effect on GLP-1-mediated satiety through stimulation of hypothalamic POMC neurons. However, db/db mice harbor a leptin receptor mutation, which causes hyperphagia, and may confound interpretation of GLP-1-regulated satiety and feeding behavior in this mouse model.

Direct evidence to support α-cell-mediated GLP-1 secretion as the mechanism of FATP2KO-associated hyperinsulinemia included (a) co-localization of FATP2 with islet α-, but not β- or δ-cells, (b) FATP2 mRNA expression in human and mouse α-, but not β-cells, (c) inhibition of fatty acid uptake by FATP2 inhibitors in α-cells, (d) increased GLP-1-positive α-cell mass in FATP2KO db/db mice, and (e) enhanced GLP-1 secretion and exendin[9-39]-inhibitable GSIS in FATP2 inhibitor-treated human islets. While these results strongly support intra-islet GLP-1 stimulation of insulin as the mechanism of FATP2KO-associated glucose reduction, we also cannot exclude the contribution of reduced plasma glucagon in FATP2KO db/db mice (Supplementary Figure 3).

Prior investigation of fatty acid effects on GLP-1 secretion is limited. Incubation of a fatty acid mixture with α-TC1-6 cells stimulated GLP-1 secretion at low glucose, and suppressed GLP- 1 in high glucose conditions (44). Similar results were observed in L-cells, with palmitate inhibition of GLP-1 secretion (46). The stimulation of GLP-1 is primarily by unsaturated fatty acids (44, 47), and is presumed to be mediated by the G-protein coupled free fatty acid receptor, FFAR4, in α-cells (48) and FATP4 or FFAR1 in L-cells (47, 49).

The mechanism by which FATP2 inhibition stimulates GLP-1 is unknown. Palmitate inhibits islet and L-cell PC1/3 (50–52), the enzyme that catalyzes the conversion of proglucagon to GLP-1, and proinsulin to insulin in β-cells (53). Although the islet studies focused on PC1/3 effects on proinsulin cleavage (50–52), it is conceivable that FATP2 inhibition also enhances GLP- 1 by blocking palmitate-induced PC1/3 suppression in α-cells. Other potential mechanisms of enhanced α-cell GLP-1 secretion include inhibition of lipotoxicity (44, 46) and reciprocal stimulation of GLP-1 by insulin. While a feed-forward (insulin stimulating GLP-1) mechanism has been proposed for intestinal L-cells (54), similar results in α-cells have not been described.

The insulinotropic effect of GLP-1 on β-cells is due primarily to GLP-1 receptor-mediated augmentation of GSIS. However, sustained hyperinsulinemia in FATP2KO db/db mice was also facilitated by increased β-cell mass and islet hypertrophy, which is a predominately GLP-1- independent process. GLP-1 exerts anti-apoptotic/cytoprotective effects of GLP-1 on β-cells (55, 56), but in human islets GLP-1 does not stimulate β-cell mitogenesis or affect islet size (56, 57). Identification of the FATP2KO-regulated factors that stimulate islet hypertrophy will require further investigation.

The effects of fatty acids on insulin secretion depend on many factors, including fatty acid carbon chain length and saturation, chronicity of exposure, fasting state, and concomitant glucose concentration (16, 58). While palmitate suppressed GLP-1 secretion in αTC1-6 cells (Figure 7C), we observed a more modest effect of palmitate on insulin secretion in human islets (Figure 8). However, even in the absence of fatty acids, GSIS was enhanced in Lipofermata-treated islets, suggesting that FATP2 inhibition may mediate insulinotropic effects independent of fatty acid uptake. To date, FATP2 has been linked primarily to lipid metabolism, particularly PPARα-regulated genes (6), which have no effect on pancreatic insulin release (59).

Downstream pathways from FATP2 are largely unexplored, and identification of additional FATP2-directed events, which may regulate GLP-1-dependent GSIS, are therefore warranted.

We conclude that FATP2 deletion or inhibition exerts glucose-lowering effects through α-cell-mediated GLP-1 secretion and paracrine β-cell insulin release. One potential clinical implication is that in contrast to diabetes treatment with GLP-1 receptor agonists, FATP2 inhibition may represent a more natural stimulus of α-cell GLP-1 augmentation. Moreover, FATP2 inhibition could represent a potential adjunctive glucose-lowering therapy, and/or a means to delay onset of type 2 diabetes.

## ACKNOWLEDGMENTS

The authors thank Hanxiao Liu and Dr. Yan Li, CWRU Dept of Genetics, for assistance with islet selection and culture methods.

## Funding

This work was supported by grants from the National Institutes of Health (5R01DK064819 to DA, 2R01DK067528 to JRS).

## Author contributions

SK, RJG, ZL, VL conducted experiments, acquired data, analyzed data, and reviewed/edited the manuscript; IS, contributed reagents, contributed to discussion and reviewed/edited the manuscript; JS contributed to discussion and reviewed/edited the manuscript; PO-O conducted experiments, acquired data and reviewed/edited the manuscript; JLG conducted experiments, analyzed data, and reviewed/edited the manuscript; DA designed research studies, analyzed data and reviewed/edited the manuscript; JRS obtained funding, designed research studies, analyzed data, wrote and edited the manuscript.

---

**Figure S1.**
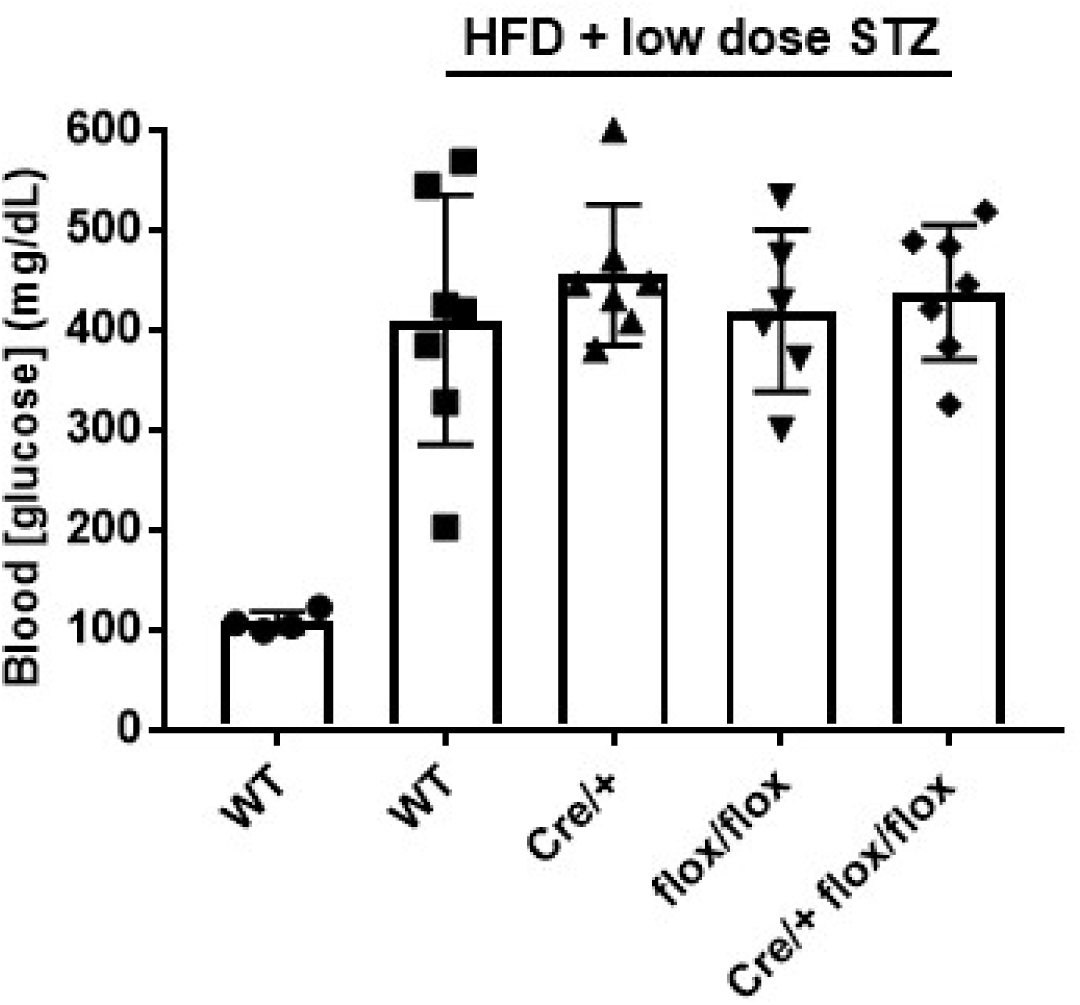
Blood glucose in mice with proximal tubule FATP2 gene deletion. Six months old C57BLKS/J wild-type (WT), Cre-lox, Cre and floxed controls with diabetes induced by high fat diet (HFD) and streptozotocin (STZ) (see Methods for details) were fasted for four hours, and tail vein blood glucose was measured by glucometer. Blood glucose was normal for all genotypes in the absence of HFD and STZ (not shown).

**Figure S2.**
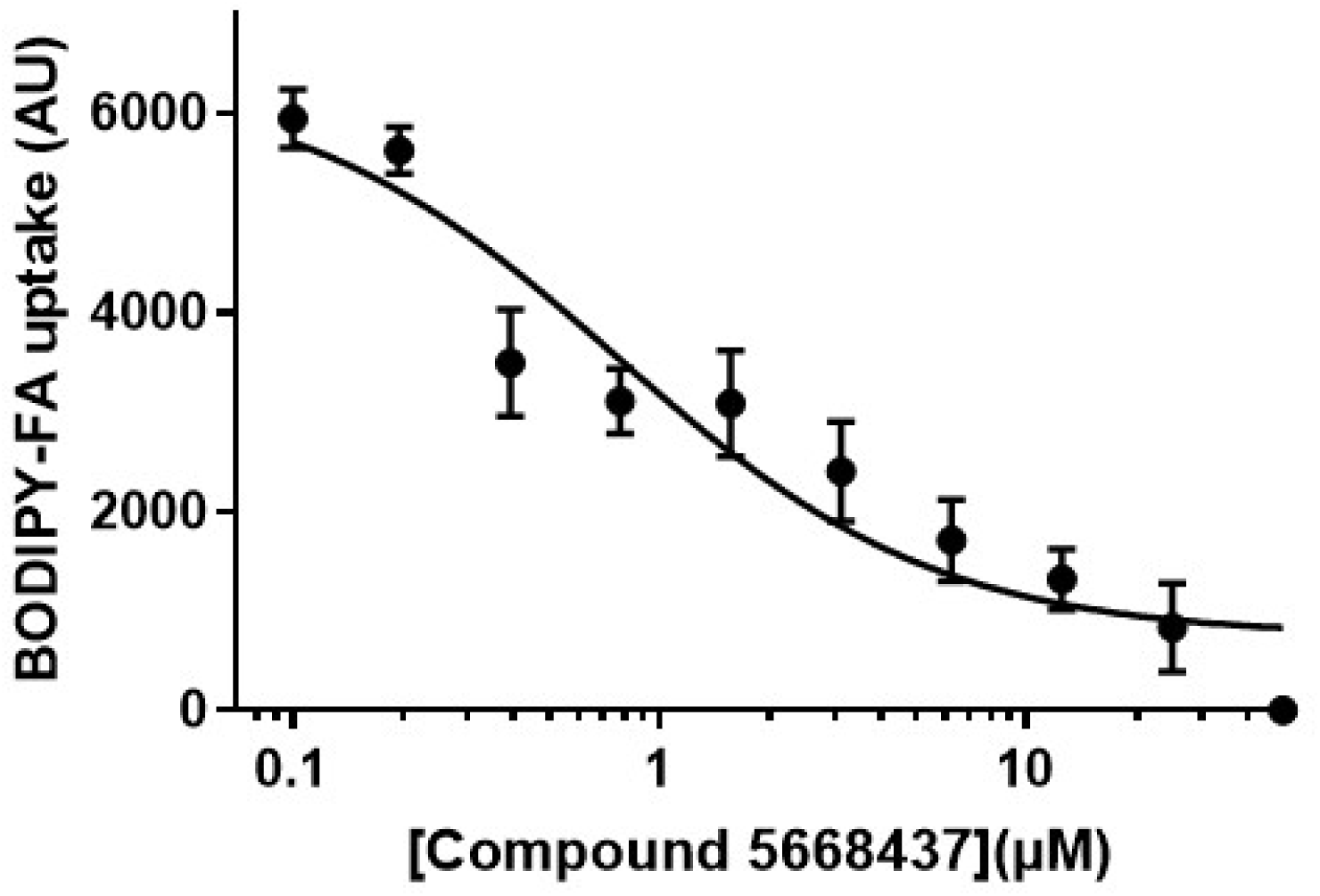
Mouse aTCl-6 cells were preincubated with compound 5668437 (1 hr, 37° C, 0-50 µM) in triplicate. BODIPY-labeled fatty acid uptake velocity was then determined, as described in Methods. Results are from four experiments.

**Figure S3.**
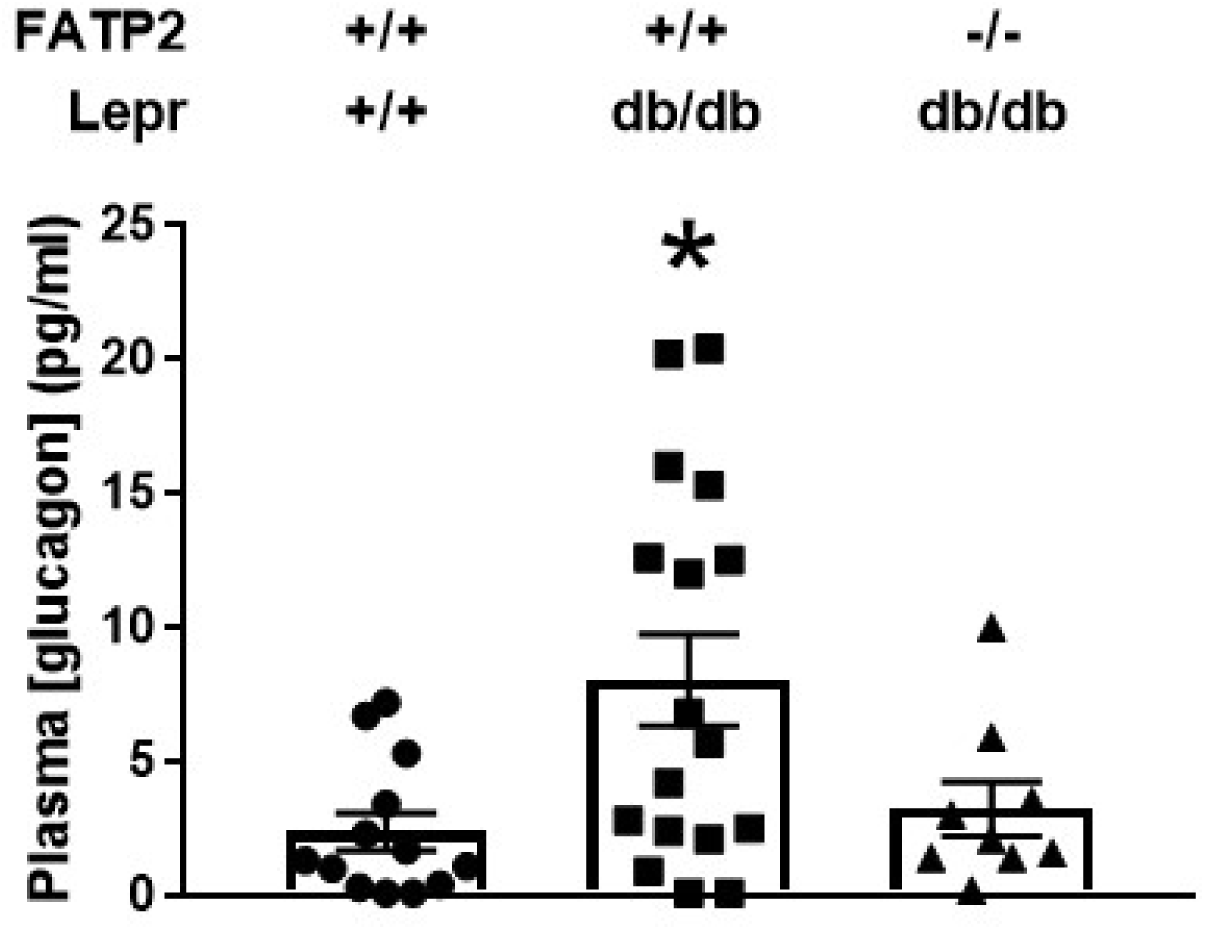
Random plasma glucagon concentrations in four to six months old wild­ type and db/db mice± FATP2 gene deletion. *P <0.05 compared to wild-type.

**Figure S4.**
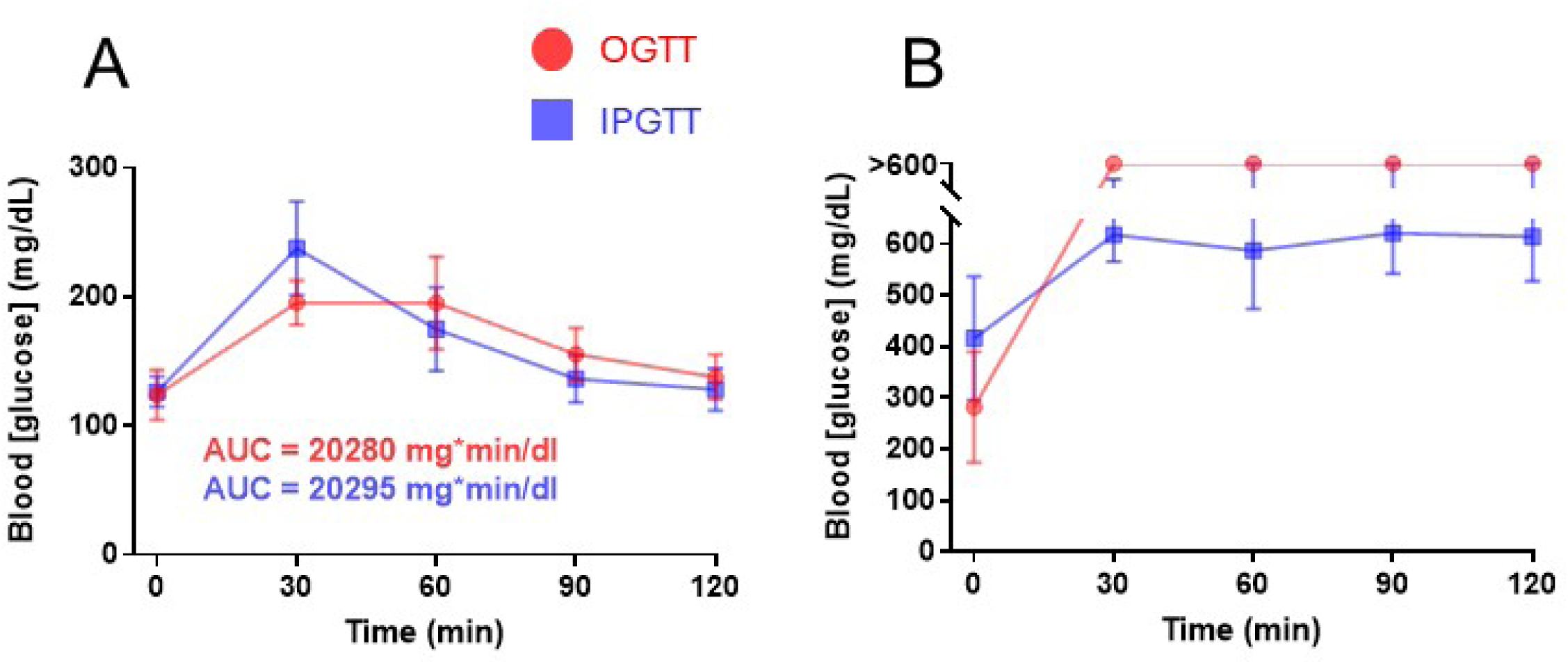
OGTT and IPGTT were conducted in FATP2KO (A) and FATP2KO db/db eNos-1- (B) mice, as described in Methods. Blood glucose was determined at the indicated times in five mice per group.

**Figure S5.**
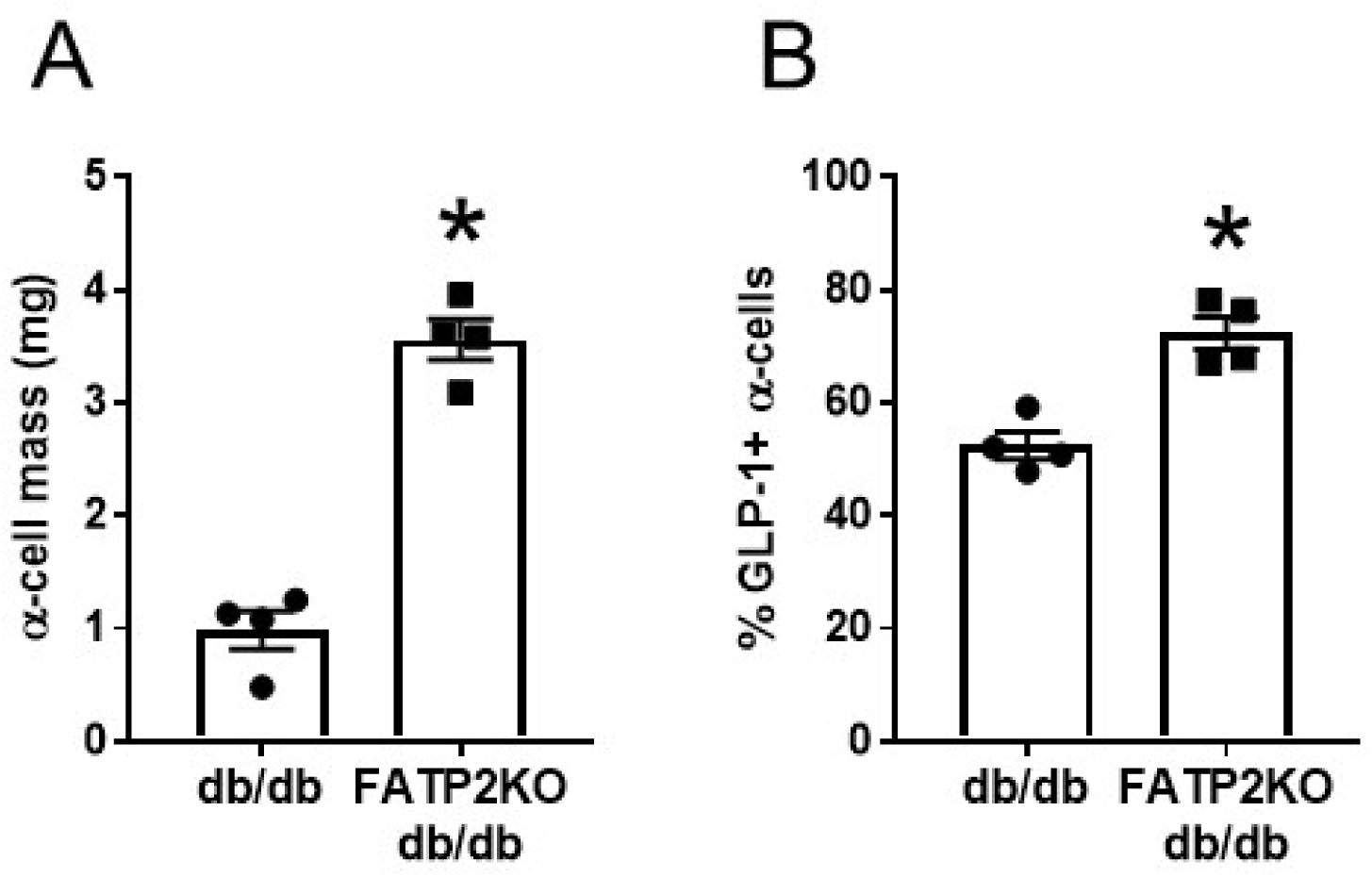
α-cell mass (A) and% GLP-1+ α­ cells (B) were determined as described in Methods in islets from db/db and FATP2KO db/db mice.* P <0.01 compared to db/db group.

**Table S1.**
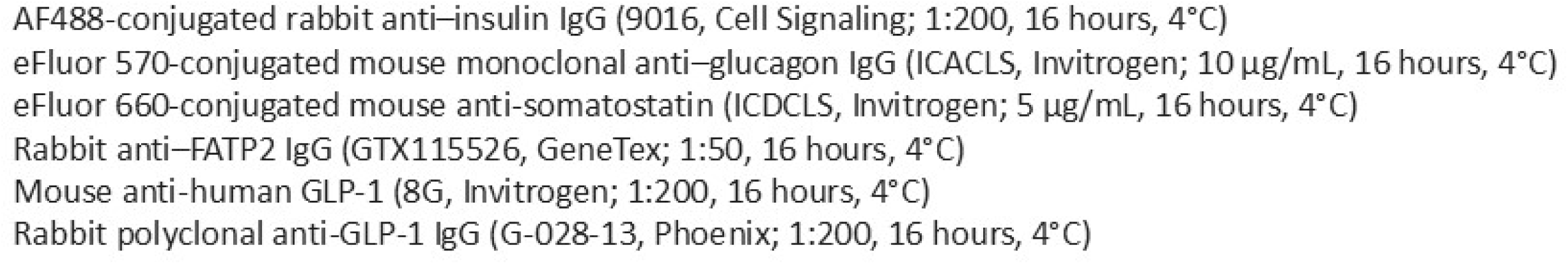
Primary antibodies and conditions.

**Table S2.**
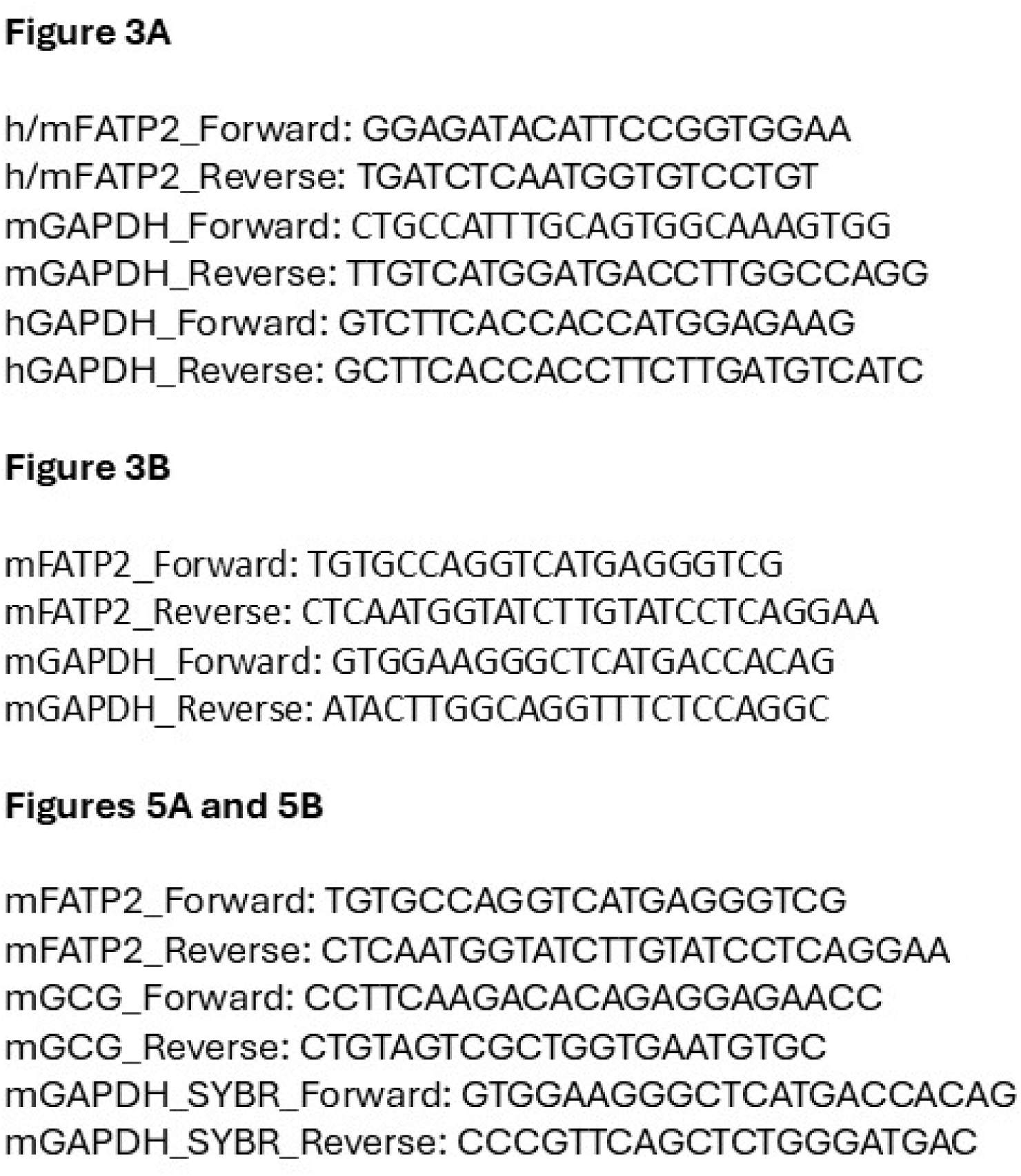
PCR primers.

## Notes

### Competing Interest Statement

The authors have declared no competing interest.

